# SERCA activity is reduced in *DJ-1* mutant flies and human cells due to oxidative modification

**DOI:** 10.1101/2020.11.19.389841

**Authors:** Cristina Solana-Manrique, Verónica Muñoz-Soriano, Francisco José Sanz, Nuria Paricio

## Abstract

*DJ-1* is a causative gene for familial Parkinson’s disease (PD) with different functions, standing out its role against oxidative stress (OS). Accordingly, PD model flies harboring a mutation in the *DJ-1β* gene (the *Drosophila* ortholog of human *DJ-1*) show high levels of OS markers like protein carbonylation, a common post-translational modification that may alter protein function. To increase our understanding of PD pathogenesis as well as to discover potential therapeutic targets for pharmacological intervention, we performed a redox proteomic assay in *DJ-1β* mutant flies. Among the proteins that showed increased carbonylation levels in PD model flies, we found SERCA, an endoplasmic reticulum Ca^2+^ channel that plays an important role in Ca^2+^ homeostasis. Several studies have supported the involvement of Ca^2+^ dyshomeostasis in PD. Interestingly, a functional link between *DJ-1* and Ca^2+^ homeostasis maintenance was previously reported. Thus, we decided to study the relation between SERCA activity and PD physiopathology. Our results showed that SERCA enzymatic activity is significantly reduced in *DJ-1β* mutant flies, probably as a consequence of OS-induced carbonylation, as well as in a human cell PD model based on *DJ-1*-deficiency. Indeed, higher carbonylation levels of SERCA were also observed in *DJ-1*-deficient SH-SY5Y neuron-like cells compared to controls. In addition, we demonstrated that SERCA activity was increased in both PD models after treatment with a specific activator of this protein, CDN1163. Consistently, CDN1163 was also able to restore PD-related phenotypes in PD model flies and to increase viability in the human cell PD model. Taken together, our results indicate that impaired SERCA activity in both familial PD models may play a role in PD physiopathology. In addition, we demonstrate that therapeutic strategies addressing SERCA activation could be beneficial to treat this disease as shown for CDN1163.

## INTRODUCTION

Parkinson’s disease (PD) is the second most common neurodegenerative disorder after Alzheimer’s disease (AD). It is pathologically characterized by the progressive loss of dopaminergic (DA) neurons in the substantia nigra of the brain, which is responsible for the typical motor dysfunction observed in PD patients (Poewe et al., 2017). Although most PD cases are sporadic, 10% of them are monogenic forms of the disease caused by mutations in specific genes (Lill, 2016). The study of their function has been important in shedding light in the pathogenic cellular events underlying PD (Poewe et al., 2017).

One of the genes involved in PD is *PARK7*, which was first linked to early onset, familial forms of PD in 2003 (Bonifati et al., 2003). It encodes the DJ-1 protein to which a range of different cellular and biochemical activities have been ascribed. These include roles as a reactive oxygen species (ROS) scavenger, cysteine protease, redox-regulated chaperone and transcriptional modulator, being also involved in anti-apoptotic signaling processes and RNA regulation (Repici & Giorgini, 2019). Despite this, how DJ-1 contributes to PD pathogenesis is still unclear (Poewe et al., 2017). Among the molecular processes underlying PD, calcium (Ca^2+^) dyshomeostasis has been widely described to contribute to the appearance and progression of the disease (Alvarez et al., 2020; Zaichick et al., 2017). The first functional link between *DJ-1* and maintenance of Ca^2+^ homeostasis was reported in Shtifman et al. (2011), who showed that loss of *DJ-1* function in primary skeletal muscle cells obtained from *DJ-1* null mice led to an increase in cytoplasmic Ca^2+^. As expected, *DJ-1* mutant cells exhibited reduced Ca^2+^ release from the endoplasmic reticulum (ER) in response to electrical stimulation (Shtifman et al., 2011). The Ca^2+^ channels RyR (Ryanodine Receptors) and SERCA (sarco/ER Ca^2+^-ATPase) are essential for maintenance of myoplasmic Ca^2+^ concentration and regulation of Ca^2+^ release (Avila et al., 2019). However, the authors demonstrated that expression levels of these proteins were not affected in *DJ-1* mutant cells, and suggested that gene products whose expression is dependent on the DJ-1 protein could modulate the function of either RyRs or SERCA (Shtifman et al., 2011). Aberrations in the finely-tuned intraneuronal Ca^2+^ homeostasis may have damaging effects, leading to the emergence of brain pathology in several neurodegenerative diseases (Alvarez et al., 2020). Indeed, Ca^2+^ handling proteins seem to play an important role in oxidative stress (OS), the generation of free radicals as well as mitochondrial dysfunction, and alterations in their function lead to ER stress (S. W. Park et al., 2010; Zaichick et al., 2017). Thus, SERCA dysfunction has been associated to ER stress and neuron loss in neurodegenerative disorders (Britzolaki et al., 2020). Consistently, it was shown that SERCA overexpression and/or pharmacological activation of this protein with small allosteric specific activators may alleviate ER stress (Kang et al., 2016; Sunderhaus et al., 2019). In fact, a small molecule activator of SERCA (CDN1163) was able to increase ER Ca^2+^ content, to rescue neurons from ER stress-induced cell death in vitro, and to efficiently alleviate dyskinesia in the rat 6-hydroxydopamine (6-OHDA) model of PD.

OS plays an important role in the development of PD (Hemmati-Dinarvand et al., 2019; Sanders & Timothy Greenamyre, 2013). Several factors contribute to the increase of ROS levels in DA neurons, such as dopamine metabolism, mitochondrial alterations or neuroinflammation, which in turn leads to oxidative modifications of proteins, lipids and nucleic acids (Blesa et al., 2015). Protein carbonylation represents the most common type of post-translational modification that proteins may suffer either by oxidative (direct) or non-oxidative (indirect) mechanisms. OS-induced carbonyl modification of proteins has several consequences either affecting their function or even promoting its elimination (Hecker & Wagner, 2018; Solana-Manrique et al., 2020; Suzuki et al., 2010). It has been shown that its effect on protein function depends on levels of carbonyl groups present in the molecule: while a high degree of carbonylation is correlated with protein dysfunction, moderate carbonylation may directly activate or inhibit the activities of target proteins as well as their selective proteasomal degradation (Hecker & Wagner, 2018). Indeed, identification of oxidized proteins in cellular and animal models of several human diseases by means of redox proteomics assays has been crucial to determine cellular pathways affected in patients (Butterfield & Dalle-donne, 2014; Ren et al., 2014; Shen et al., 2015; Sultana et al., 2013). Few redox proteomics studies have been performed so far in PD models (Bulteau et al., 2017; Chiaradia et al., 2019; Di Domenico et al., 2012; Poon et al., 2005), despite they could be useful to increase our understanding of PD pathogenesis as well as to discover potential therapeutic targets for pharmacological intervention.

In this scenario, the present work seeks to identify the target proteins of OS in a *Drosophila* PD model based on inactivation of *DJ-1β*, the fly ortholog of human *DJ-1* (Lavara-Culebras & Paricio, 2007; Park et al., 2005). It was previously shown that 15-day-old *DJ-1β* mutants exhibited increased ROS and protein carbonylation levels when compared to control flies of the same age (Casani et al., 2013; Lavara-Culebras et al., 2010). To achieve this, we performed redox proteomics analyses in PD model and control flies, being able to identify proteins differently carbonylated in *DJ-1β* mutants that could be involved in several cellular processes associated with the disease. Among those that exhibited increased carbonylation levels in PD model flies we found SERCA, a protein encoded by the *Drosophila Ca-P60A* gene (Sanyal et al., 2005) also known as *dSERCA*. Our results demonstrate that SERCA enzymatic activity is significantly reduced in *DJ-1β* mutant flies as well as in *DJ-1-*deficient SH-SY5Y neuron-like cells compared to controls. These results indicate that SERCA carbonylation leads to its inactivation in those PD models, as found in mammals (Žižková et al., 2014, 2018). Furthermore, we show that activation of SERCA with a specific allosteric activator of this protein (CDN1163) is capable to suppress PD-related phenotypes in fly and human cell models based on *DJ-1* deficiency. Therefore, our results support the crucial role of SERCA activity in PD pathology, and demonstrate that targeting SERCA activation is a promising strategy to develop disease-modifying therapeutics for PD.

## MATERIALS AND METHODS

### *Drosophila* strains

The stocks used in this work were: *y,w* (<u>B</u>loomington <u>D</u>rosophila <u>S</u>tock <u>C</u>enter #6598*: y*^*1*^,*w*^*1118*^), *w; DJ-1β*^*ex54*^ (hereafter called *DJ-1β*; Park et al. 2005), *arm-*GAL4 (BDSC #1560: *P{GAL4-arm.S}11*), UAS*-iRdSERCA* (Vienna Drosophila Research Center #3017: *w*^*1118*^; *P{GD1508}v3017*). All stocks and crosses were cultured on standard *Drosophila* food at 25°C, unless otherwise indicated. The *y,w* line was used as wild-type control in experiments with the *DJ-1β* strain. For RNAi experiments, the *arm-*GAL4 driver line was crossed with UAS-*iRdSERCA* to ubiquitously downregulate *dSERCA* expression. Progeny of this cross is referred to as *iRdSERCA*. In these experiment, progeny of crosses between *y,w* and *arm-GAL4* flies was used as control.

### Redox proteomics assay

#### Carbonylated protein derivatization

Groups of fifty 15-day-old female flies were homogenized in extraction buffer [8 M urea, 25 mM Tris pH 8, Halt Protease Inhibitor Cocktail (Thermo Scientific)] with a steel bead in a TissueLyser LT (Qiagen) for 5 min at 50 Hz at 4°C. Fly extracts were centrifuged three times at 9000g for 10 min at 4°C. After centrifugation, supernatant was collected and proteins were quantified using a NanoDrop 2000 spectrophotometer (Thermo Scientific). For protein precipitation, the 2-D Clean-Up Kit (GE Healthcare) was used following manufacturer’s instructions. Subsequently, 300 μg of protein were derivatized using the BodipyFL hydrazide (Invitrogen) as described in Tamarit et al. (2012).

#### 2D immunoblotting

Derivatized-protein samples were analysed by two-dimensional (2D) gel electrophoresis using Immobiline^®^ DryStrips pH 3-11 (IPG) non-linear (NL) 24 cm (GE Healthcare) for isoelectric focusing (IEF) as the first dimension and SDS-PAGE separation as the second dimension. Samples were supplemented with 65 mM DTT and 1% ampholytes (Healthcare GE). IPG strips were rehydrated overnight with DeStreak™ Rehydration Solution (GE Healthcare) and 1% (v/v) ampholytes (pH 3-11 NL; GE Healthcare) following manufacturer’s instructions. IEF conditions were performed with the following voltage program: 300 V for 4 h, linear ramp to 1000 V over 6 h, linear ramp to 8000 V over 3 h, then 8000 constant for a total focusing time of 28000 Vh at 20°C. After IEF, the IPG strips were reduced for 15 min with 2% (w/v) DTT in 50 mM Tris, 6 M urea, 30% (v/v) glycerol, 2% (w/v) SDS, and subsequently for 15 min in the same buffer containing 2.5% (w/v) iodoacetamine instead of DTT. In the second dimension, proteins were separated on 12.5% (v/v) acrylamide gels of 25 cm x 21 cm x 1 mm using an Ettan DALTsix electrophoresis unit (GE Healthcare). The SDS-PAGE was first run at a current of 2 W/gel for 1 h and then at a constant current of 15 W/gel for 5 h. After gel electrophoresis, total protein content in the gels was stained with SYPRO^®^ Ruby Protein Stain (Invitrogen) following manufacturer’s instructions. Stained gels were imaged using a TyphoonTM 9400 Variable Mode Imager (GE Healthcare) using the filter that adjusted to an emission wavelength of 510 nm and 610 nm for the BODIPY dye and for the SYPRO^®^ Ruby dye, respectively. Samespots software version 5.0 (TotalLab) was used to compare protein carbonylation between control and *DJ-1β* flies. Proteins equally expressed in both genotypes whose carbonylation levels were significantly increased (P<0.05) in PD model flies were selected for subsequent identification.

#### Mass spectrometry

Gels were silver stained by using the PlusOne Silver Staining Kit (protein, GE Healthcare) to excise the selected spots following manufacturer’s instructions. The spots were digested with sequencing grade trypsin (Promega) as described (Shevchenko et al., 1996). Digestion was stopped with TFA (1% (v/v) final concentration) and the digested peptides were concentrated to 7 μL. A BSA plug was analysed in the same way to control the digestion process. The resulting mixtures were analyzed by tandem mass spectrometry (MS/MS) in a 5800 MALDI TOFTOF (ABSciex) in positive reflectron mode (3000 shots every position). Five of the most intense precursors (according to the threshold criteria: minimum signal-to-noise: 10, minimum cluster area: 500, maximum precursor gap: 200 ppm, maximum fraction gap: 4) were selected for every position for the MS/MS analysis. MS/MS data was acquired using the default 1kV MS/MS method. The MS/MS information was sent to MASCOT via the Protein Pilot (ABSciex). Database search was performed on Unigen database. Searches were done with tryptic specificity allowing one missed cleavage and a tolerance on the mass measurement of 100 ppm in MS mode and 0.8 Da in MSMS mode. Carbamidomethylation of Cys was used as a fixed modification and oxidation of Met and deamidation of Asn and Gln as variable modifications.

For the spots that could not be identified by MS/MS, a liquid chromatography and tandem mass spectrometry (LC-MS/MS) analysis was performed. 5 μl of every sample (except the main bands) were loaded onto a trap column (NanoLC Column, 3μm C18-CL, 350 μm x 0.5 mm; Eksigen) and desalted with 0.1% (v/v) TFA at 3μl/min during 5 min. The peptides were then loaded onto an analytical column (LC Column, 3 μm C18-CL, 75 μm x 12 cm, Nikkyo) equilibrated in 5% (v/v) acetonitrile 0.1% (v/v) formic acid. Elution was carried out with a linear gradient of 5-45% B in A for 15min (A: 0.1% (v/v) formic acid; B: 0.1% (v/v) formic acid in acetonitrile) at a flow rate of 300 nl/min. Peptides were analysed in a mass spectrometer nanoESI qQTOF (5600 TripleTOF, ABSCIEX).

The tripleTOF was operated in information-dependent acquisition mode, in which a 0.25-s TOF MS scan from 350–1250 m/z was performed, followed by 0.05-s product ion scans from 100–1500 m/z on the 50 most intense 2-5 charged ions. The LC-MS/MS information was sent to ProteinPilot v4.5 search engine (ABSciex). Data obtained from the sample were analysed combined for database search. ProteinPilot default parameters were used to generate peak list directly from 5600 TripleTof .wiff files. The Paragon algorithm of ProteinPilot was used to search NCBI protein database with the following parameters: trypsin specificity, iodoacetamide cys-alkylation, taxonomy restricted to *Drosophila melanogaster*, and the search effort set to rapid. To avoid using the same spectral evidence in more than one protein, the identified proteins were grouped based on MS/MS spectra by the Protein-Pilot Progroup algorithm. Thus, proteins sharing MS/MS spectra were grouped, regardless of the peptide sequence assigned. The protein within each group that can explain more spectral data with confidence was shown as the primary protein of the group.

### Climbing assays

Locomotor ability of flies was analyzed by means of climbing assays as previously described in Sanz et al. (2017). The experiments were carried out three times and the climbing ability of mutant flies and their corresponding controls was determined as the average of the height reached by each fly after 10 s.

### Cell culture and drug treatment

SH-SY5Y neuroblastoma control cells (*pLKO.1*) and *DJ-1-*deficient cells (Sanz et al., 2017) were maintained in a selective growth medium consisting of Dulbecco’s Modified Eagle Medium/Nutrient Mixture F-12 (DMEM/F-12) (Labclinics) supplemented with 2 μg/ml puromycin (Labclinics), 10% (v/v) fetal bovine serum (FBS) (Labclinics) and 100 mg/ml penicillin/streptomycin (Labclinics) at 37 °C and 5% CO_2_.

Cell viability assays in *DJ-1*-deficient cells treated with different concentrations of CDN1163 (Sigma-Aldrich) were performed with MTT (3-(4, 5-dimethylthiazol-2-yl)-2-5-diphenyltetrazolium bromide) (Sigma-Aldrich) as described in Sanz et al. (2017), with slight modifications. Briefly, *DJ-1-*deficient cells were seeded in a 96-well plate at a density of 1.8×10^4^ cells/well and incubated for 24 h with different concentrations of CDN1163 in the range 1-50 μM, or with 0.1% dimethyl sulfoxide (DMSO) as vehicle. Subsequently they were incubated with 100 μM H_2_O_2_ for 3 h.

### SERCA activity assay

To obtain protein extracts from adult flies of different genotypes, 20 female flies were homogenized in Homogenization buffer [250 mM sucrose, 5 mM HEPES pH 7, 1 mM PMSF, Halt Protease Inhibitor Cocktail (Thermo Scientific)] with a steel bead in a TissueLyser LT (Qiagen) for 5 min at 50 Hz. Fly extracts were centrifuged at 9000*g* for 8 min at 4°C. After centrifugation supernatant was collected. Protein extracts from *DJ-1-*deficient and control SH-SY5Y cells were obtained from aliquots of 3×10^6^ cells. To do this, cells were lifted with trypsin and harvested in complete DMEM-F12 medium, centrifuged at 300g for 5 min, washed with 1% PBS buffer at 4 °C and resuspended in Homogenization buffer [250 mM sucrose, 5 mM HEPES pH 7, 1 mM PMSF, Halt Protease Inhibitor Cocktail (Thermo Scientific)]. Cell suspensions were subjected to three successive cycles of freezing in liquid N_2_, thawing on ice and vortexing. Lysates were centrifuged at 15.000g for 10 min at 4 °C and supernatant was collected. Supernatants containing fly and cell lysates were quantified with Pierce™ BCA Protein Assay Kit (Thermo Scientific) following manufacturer’s instructions. SERCA activity was measured by spectrophotometric assay using an enzyme-coupled system as described in Moraru et al. (2017). Previously, the SERCA activity assay was validated in *iRdSERCA* mutant flies, in which the expression of the gene encoding this protein is reduced. All the assays were performed in triplicate.

### RT-qPCR analyses

Total RNA from ten 15-day-old *DJ-1β*, *iRdSERCA* or control flies (*y,w* or *arm*-GAL4/+, depending on the experiment) and from aliquots of 3×10^6^ *DJ-1* mutant and control cells was extracted and reverse transcribed as described in Solana-Manrique et al. (2020).

For the RT-qPCR reactions, the following pairs of primers were used: *Drosophila dSERCA* direct primer (5’-3’); *Drosophila dSERCA* reverse primer (5’-3’); *Drosophila Tubulin* direct primer (5’-GTATCTCTATCCATGTTGGTCAGG-3’); *Drosophila Tubulin* reverse primer (5’-AGACGGCATCTG GCCATCG-3’); *Homo sapiens ATP2A1* direct primer (5’-3’); *Homo sapiens ATP2A1* reverse primer (5’-3’); *Homo sapiens ATP2A2* direct primer (5’-3’); *Homo sapiens ATP2A2* reverse primer (5’-3’); *Homo sapiens Tubulin* direct primer (5’-GCCGAGATCACCAATGCCT-3’); *Homo sapiens Tubulin* reverse primer (5’-TCACACTTGACCATTTGATTGGC-3’).

### SERCA immunoprecipitation and post-Western blot derivatizarion

To confirm the redox proteomic assay results, carbonylation levels of SERCA were detected by immunoprecipitation and post-Western blot derivatization in control and *DJ-1*-deficient cells. For protein extraction, aliquots of 3×10^6^ cells of mutant and control cells were homogenized in cell lysis buffer [25 mM Tris pH8, 0.05% (w/v) SDS, 75 mM NaCl, 0.5% (v/v) Triton X-100, 0.25% (w/v) sodium deoxycholate, 100 mM NaF, 20 mM Na_4_P_2_O_7_, 1 mM PMSF, Halt Protease Inhibitor Cocktail (Thermo Scientific)] with brief sonication. Extracts were centrifuged at 15.000g for 10 min at 4°C and supernatants were collected and quantified with Pierce™ BCA Protein Assay Kit (Thermo Scientific) following manufacturer’s instructions. Protein samples (200 μg) were incubated with anti-SERCA antibody (CaF2-5D2, Developmental Studies Hybridoma Bank) for 1 h at 4 °C with gentle shaking. Pierce™ Protein A Agarose (Thermo Scientific) beads were added, and the mixture was incubated for 4 h at 4 °C with gentle shaking. After incubation, beads were centrifuged and washed three times with cells lysis buffer. Then, beads were resuspended in SDS loading buffer [62.5 mM Tris pH 6.8, 2% (w/v) SDS, 10% (v/v) glycerol, 50 mM DTT, 0.01% (p/v) bromophenol blue] and boiled for 5 min. Proteins bound to beads were separated by SDS-PAGE and transferred to nitrocellulose membranes, which were subsequently derivatized as described in Feng et al. (2017). Carbonylation levels were detected by incubating the membranes with anti-DNP (1:250, Sigma-Aldrich) overnight at 4°C. Blots were then washed and incubated with the goat anti-rabbit IgG/HRP conjugate (1:5000, Sigma-Aldrich). The signal was detected with a Pierce™ ECL detection kit (Thermo Scientific) and captured using ImageQuant LAS4000. Images were analyzed with ImageJ software (NIH).

### CDN1163 treatments in *Drosophila*

Flies were cultured on standard *Drosophila* food containing 0.1% DMSO for untreated control experiments, or in the same medium supplemented with a final concentration of 50 μM CDN1163. Ten males and 20 females of *y,w* or *DJ-1β* flies were crossed in each vial with or without compound and maintained at 25 °C. For climbing assays, freshly eclosed flies were transferred to new vials with the compounds and the climbing assay was carried out 5 days after eclosion. For enzymatic assays, freshly eclosed flies were transferred to new vials with or without the compounds every two days for 15 days. After this time, 15-day-old flies were frozen in liquid N_2_ and stored at −80 °C.

### Quantification of protein carbonyl group formation and H2O2 levels

Protein carbonyl groups were measured in 5-day-old fly extracts using 2,4-dinitrophenylhydrazine (DNPH) derivatization in 96-well plates (Greiner 96 well plate, polypropylene) as described in Sanz et al. (2017). H_2_O_2_ levels were measured using the *Amplex H_2_O_2_ Red Kit* (Invitrogen) in 5-day-old fly extracts with as described in Sanz et al., (2017). All experiments were carried out using three biological replicates and three technical replicates for each sample.

### Statistical analysis

In all cases, data are expressed as means ± standard deviation (s.d.). The significance of differences between means was assessed using t-test. Differences were considered significant when \**P* < 0.05.

## RESULTS

### Carbonylation protein pattern in *DJ-1β* mutant flies

Previous studies performed in 15-day-old *DJ-1β* mutants showed that protein carbonylation levels were significantly increased in such flies when compared to controls of the same age (Casani et al., 2013). In order to identify specific proteins that were most susceptible to oxidative damage in PD model flies, we undertook redox proteomic analyses in 15-day-old *DJ-1β* mutants and *y,w* control flies. To our knowledge, this is the first redox proteomics assay performed in a *Drosophila* PD model. Figure 1 shows representative 2D Sypro Ruby-stained gels for total protein content (Fig. 1A) and the corresponding Bodipy staining (Fig. 1B) for oxidized proteins obtained in these experiments. The redox proteomics analyses led to identify 53 spots corresponding to differently carbonylated proteins with equivalent expression levels. Specifically, 31 of them showed increased levels and 22 showed reduced levels in 15-day-old *DJ-1β* mutants compared to control flies of the same age. Some of the differentially carbonylated proteins, evidenced by Bodipy staining, were subsequently characterized by proteomic procedures (see Materials and Methods section for details). We specifically focused on those spots corresponding to proteins with increased carbonylation levels in PD model flies, from which twelve were selected for mass spectrometry (MS) identification (see Materials and Methods). Table 1 shows the proteins identified in these experiments, the genes encoding them and the fold increase in oxidation of each protein in *DJ-1β* mutants compared to control flies, which ranges from 1.4 to 2.8. The functions of the identified proteins are diverse (see Table S1). Two of them, enolase (Eno) and phosphofructokinase (Pfk), are glycolytic enzymes and were shown to present enhanced activity in PD model flies compared to controls (Solana-Manrique et al., 2020). Other proteins identified were: voltage-dependent anion-selective channel (VDAC), actin, vitellogenin-2 and 3, tyrosyl-tRNA synthethase, myosin heavy chain, V-type proton ATPase catalytic subunit A isoform 2, arginine kinase and citrate synthase. Interestingly, this is the first study in which some of these proteins are shown to be specifically carbonylated in an animal model of familial PD.

**Figure 1.**
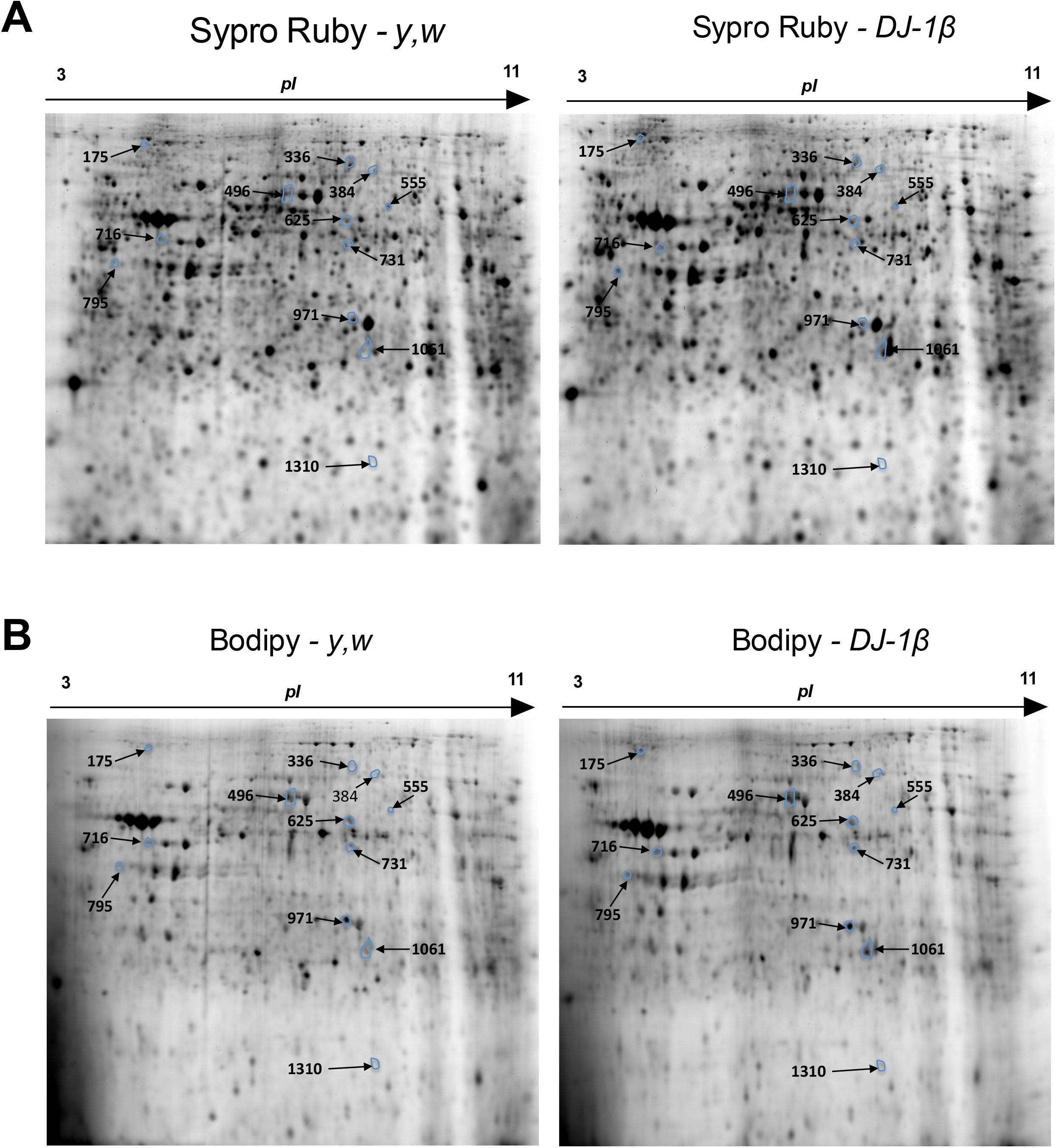
Representative 2D gels obtained in the redox proteomics assays from 15-day-old control (*y,w*) and *DJ-1β* mutant flies. (A) Representative 2D gels stained with Sypro Ruby to visualize total protein content and with (B) Bodipy to reveal carbonyl groups in whole protein extracts of 15-day-old control (*y,w*) and *DJ-1β* mutant flies. The spots showing increased carbonylation levels in PD model flies compared to controls and selected for protein identification are indicated with arrows. The spot numbers are the same as those listed in Table 1.

**Table 1.**
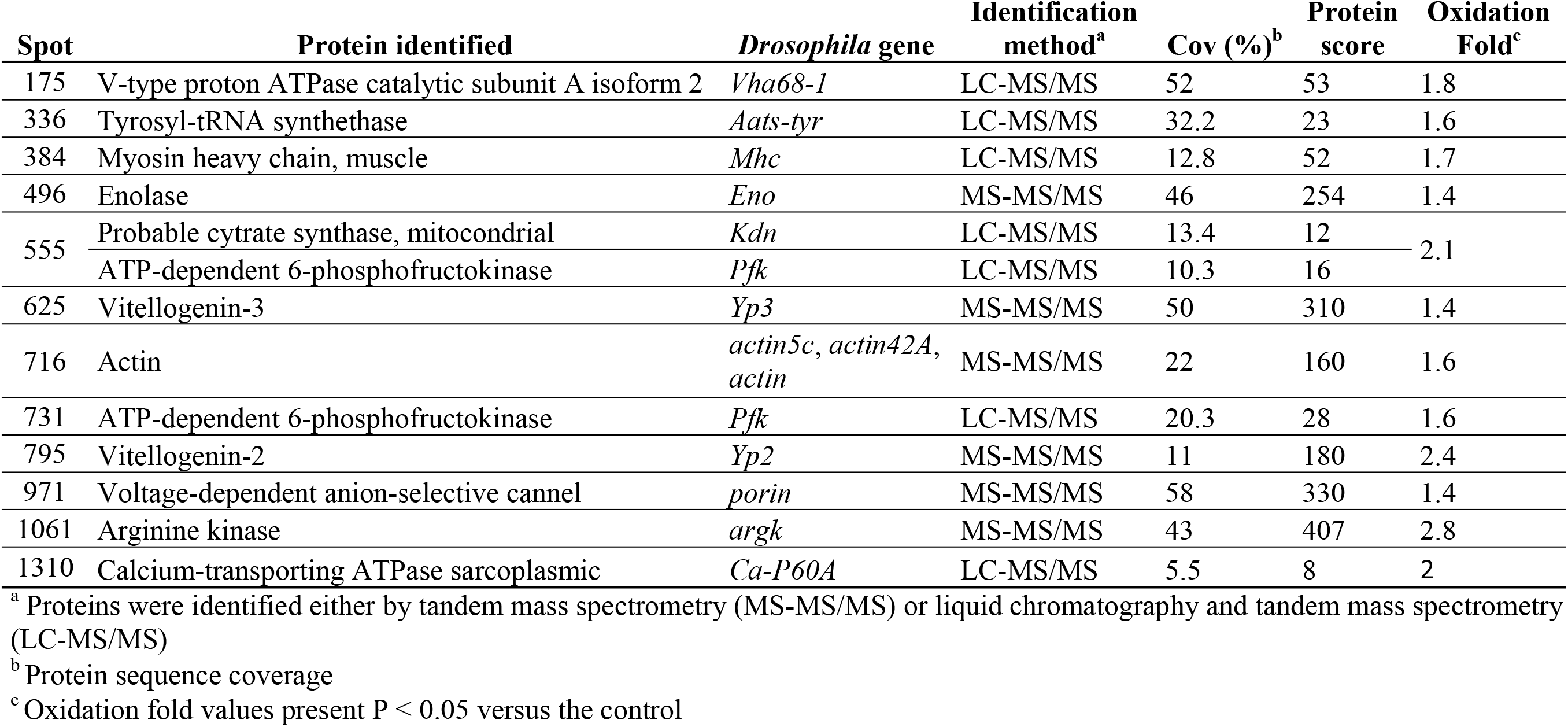
Proteins identified by mass spectrometry that showed increased carbonylation levels in 15-day-old *DJ-1β* mutant flies.

Among the proteins identified, SERCA (EC 7.2.2.10) is an ion channel associated to Ca^2+^ homeostasis and located in the ER. It balances calcium between the ER and the cytosol by pumping Ca^2+^ into the ER in order to ensure proper protein folding and chaperone function (Britzolaki et al., 2020, Sunderhaus et al., 2019). Although its exact role in the central nervous system is not clear yet, it is known that SERCA-mediated Ca^2+^ dyshomeostasis is associated with neuropathological conditions (Alvarez et al., 2020; Britzolaki et al., 2020). Several studies indicate that inhibition of SERCA function is related to neurodegeneration (Britzolaki et al., 2020; Sunderhaus et al., 2019), thus suggesting that this could be a major cause of PD (Alvarez et al., 2020). Therefore, we suspected that alterations of SERCA function caused by high carbonylation levels could be contributing to PD-related phenotypes in *DJ-1β* mutant flies as well as to PD physiopathology by affecting Ca^2+^ homeostasis.

### SERCA activity is reduced in *DJ-1β* mutants

As mentioned above, one of the proteins that exhibited increased carbonylation levels in 15-day-old *DJ-1β* mutants compared to control flies of the same age was the SERCA Ca^2+^ pump. To determine the consequences of such posttranslational modification on SERCA function, we decided to measure its activity in *DJ-1β* mutants by using a protocol already tested in *Drosophila* (Moraru et al., 2017). We first validated the SERCA activity assay in protein extracts from *iRdSERCA* flies, in which the expression of the *dSERCA* gene was ubiquitously downregulated with the *arm*-GAL4 driver. As expected, SERCA activity was significantly reduced in mutant flies compared to controls (Fig. 2A), due to a decrease in *dSERCA* expression levels in such flies (Fig. 2B). Subsequently, we analyzed SERCA activity in 15-day-old *DJ-1β* mutants and *y,w* control flies using the same assay. Our results showed that SERCA activity was significantly reduced in PD model flies compared to controls (Fig. 2A), despite that *dSERCA* expression levels were equivalent in both fly strains (Fig. 2B). Consistently, we found that 15-day-old *iRdSERCA* mutants showed phenotypes similar to those exhibited by *DJ-1β* mutants (Lavara-Culebras & Paricio, 2007). Hence, *iRdSERCA* mutant flies displayed motor impairments, reflected by a reduced climbing ability, and also a decreased lifespan when compared to control flies (Fig. S1). These results are in agreement with previous reports in which other *dSERCA* mutants were analyzed (Kaneko et al., 2014; Sanyal et al., 2005). Taken together, our results support the existence of a functional link between *DJ-1* and Ca^2+^ homeostasis maintenance, as previously suggested in mice (Shtifman et al., 2011), and indicate that loss of *DJ-1β* function may have a direct impact on the activity of this Ca^2+^ pump probably due to high ROS levels.

**Figure 2.**
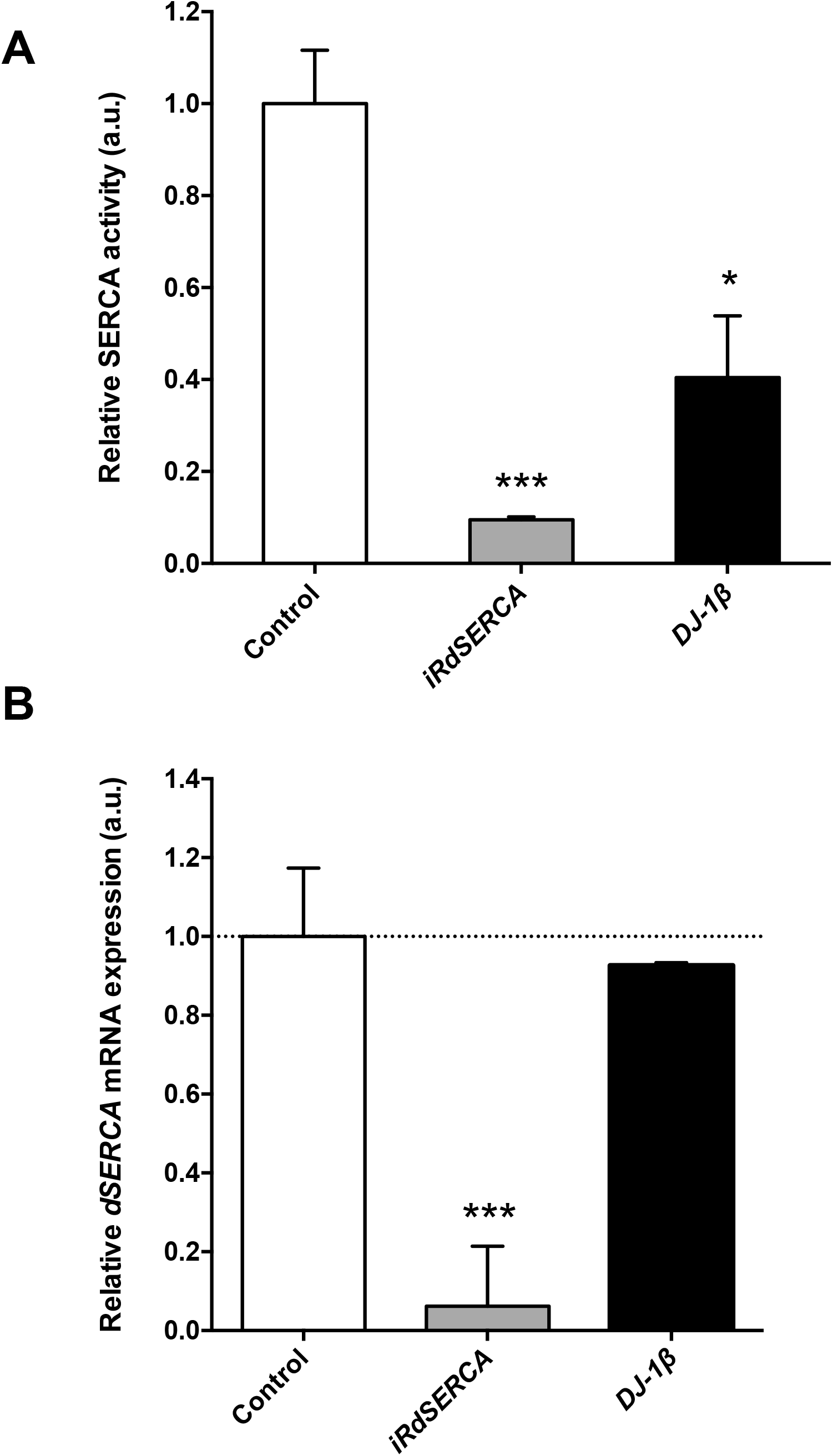
Analysis of SERCA activity in *DJ-1β* mutant flies. (A) SERCA activity was measured in 15-day-old *y,w* control flies, and in *iRdSERCA* and *DJ-1β* mutant flies. In all cases results were relativized to data obtained from 15-day-old *y,w* control flies and are expressed as arbitrary units (a.u.). (B) Analysis of *dSERCA* gene expression levels by RT-qPCR analysis. *dSERCA* expression levels were measured in 15-day-old *DJ-1β* mutants and *iRdSERCA* flies. Results are relativized to data obtained in control flies for each genotype (*y,w* and arm-GAL4/+ flies, respectively), and are expressed as arbitrary units (a.u.). For (A) error bars show s.d. from three independent experiments in which three biological replicates were used. For (B) error bars show s.d. from four independent experiments (*, *P* < 0.05; ***, *P* < 0.001).

### SERCA activity is reduced in *DJ-1*-deficient cells due to increased carbonylation levels

To confirm the results obtained in *Drosophila*, we analyzed SERCA activity in an *in vitro* human cell PD model based on *DJ-1*-deficiency (Sanz et al., 2017). We and others already showed that those cells were more susceptible than control cells to OS-induced cell death (Gao et al. 2011; Sanz et al. 2017; Solana-Manrique et al. 2020; Wang et al. 2011). Therefore, SERCA activity was determined under OS conditions induced with 50 μM H_2_O_2_, a concentration that significantly decreased viability of *DJ-1*-deficient cells compared to controls but in which enough cells could be harvested (Solana-Manrique et al., 2020). Our results showed that SERCA activity was significantly reduced in *DJ-1*-deficient cells (Fig. 3A). To exclude the possibility that variations in SERCA activity between *DJ-1*-deficient and control cells were reflecting differences in gene expression levels, we performed RT-qPCR assays. Using the sequence of the *Drosophila dSERCA* gene (CG3725) as a query, we found different *SERCA* orthologs in humans. Among them, we selected those genes that are significantly expressed in SH-SY5Y cells according to the Human Protein Atlas (Uhlén et al., 2010, 2015) such as *ATP2A1* and *ATP2A2*, which encode the SERCA1 and SERCA2 isoforms, respectively. No significant differences were detected in *ATP2A1* and *ATP2A2* expression levels between *DJ-1*-deficient and control cells (Fig. 3B). These results are consistent with those found in skeletal muscle of *DJ-1* null mice in which no significant alterations in the steady-state expression levels of the SERCA1 protein were found (Shtifman et al., 2011). To determine whether reduction of SERCA activity could be associated to its oxidative modification, we decided to analyze SERCA carbonylation levels in PD model and control cells. To do so, we performed SERCA immunoprecipitation with an anti-SERCA1 antibody and post-Western blot derivatization of protein extracts using an anti-DNP antibody. As shown in Fig. 3C, the SERCA protein was equally expressed in both cell lines but showed significantly increased carbonylation levels in *DJ-1*-deficient cells. These results confirmed that SERCA activity was reduced in the human cell PD model based on *DJ-1* deficiency due to an increase in its carbonylation levels, and validated the results obtained in the redox proteomics assays with *DJ-1β* mutant flies.

**Figure 3.**
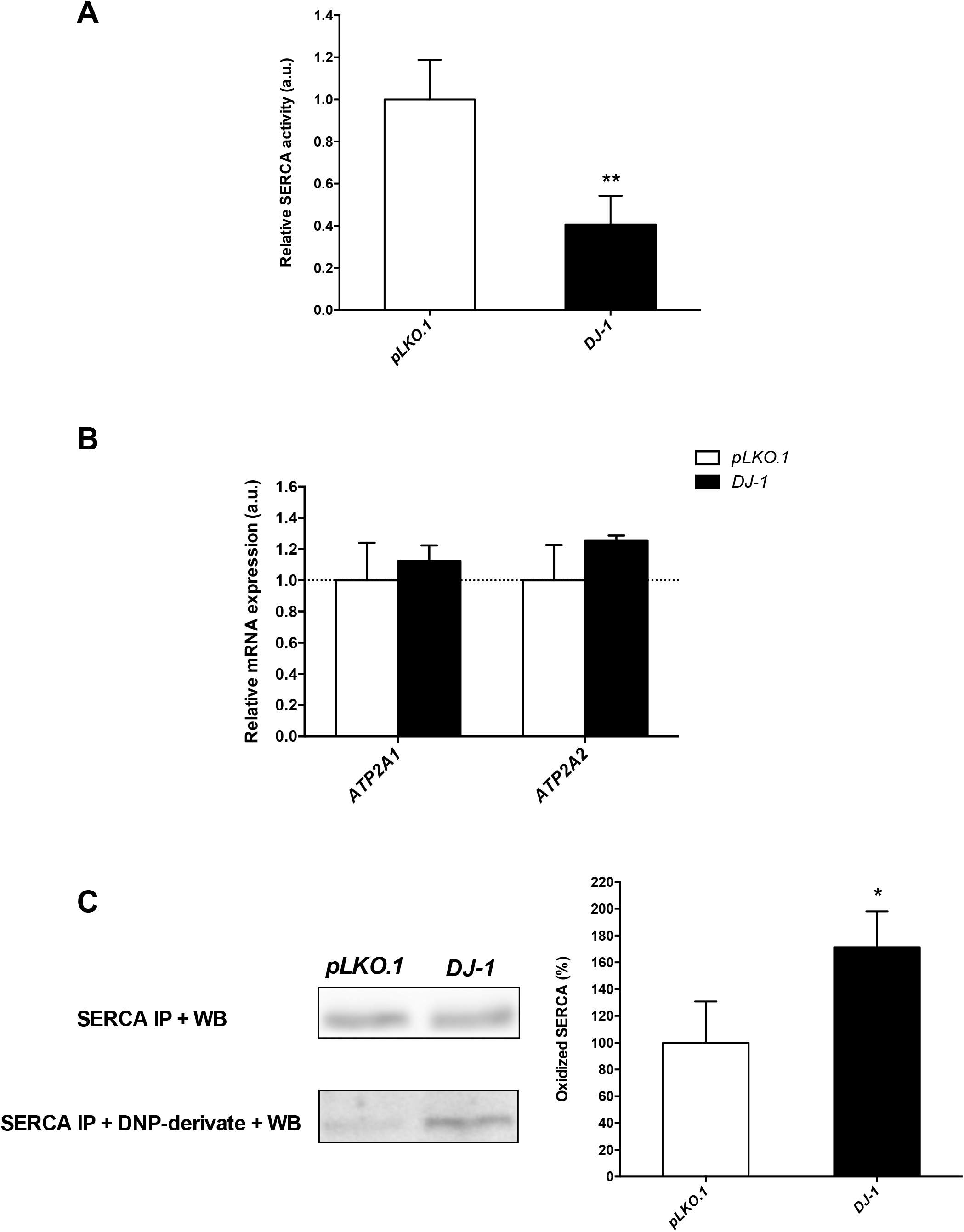
Analysis of SERCA activity and carbonylation levels in *DJ-1*-deficient human cells. (A) SERCA activity was measured in *DJ-1*-deficient cells under OS conditions induced with 50 μM H_2_O_2_. (B) Graphical representation of *ATP2A1* (SERCA1) and *ATP2A2* (SERCA2) expression levels in *DJ-1*-deficient cells under OS condition induced with 50 μM H_2_O_2_. Results are referred to data obtained in *pLKO.1* control cells under the same OS condition, and are expressed as arbitrary units (a.u.). In both cases, *tubulin* expression levels were measured and used as an internal control for RNA amount in each sample. (C) SERCA carbonylation levels were detected by inmunoprecipitation and post-Western blot analysis in *DJ-1*-deficient cells under OS conditions induced with 50 μM H_2_O_2_. Left panel shows representative western blots and the right panel constitutes its graphical representation. Results were relativized to data obtained in *pLKO.1* control cells also treated with 50 μM H_2_O_2_, and are expressed as arbitrary units (a.u). Error bars show s.d. from three replicates and three independent experiments in A and C, and from four independent experiments in B (*, *P* < 0.05; ***, *P* < 0.001).

### The SERCA activator CDN1163 suppresses phenotypes in *DJ-1β* mutant flies and *DJ-1-* deficient cells

Dysregulation of Ca^2+^ homeostasis and ER stress have been linked to neurodegeneration and to the development of neurodegenerative diseases, such as PD (Britzolaki et al., 2020; Dahl, 2017; Rahate et al., 2020). The SERCA pump is a protein whose main function is to transport Ca^2+^ into the ER, hence maintaining physiological cytoplasmic Ca^2+^ levels, and its activation has been associated to an alleviation of ER stress (Dahl, 2017; Qaisar et al., 2019). Therefore, we decided to test if an increase of SERCA activity could attenuate phenotypes in our fly and human cell models of PD by using the compound CDN1163 (Dahl, 2017). CDN1163 is a quinolone-amide that acts as an allosteric activator of SERCA, directly binding to this protein. First, we tested the effect of CDN1163 in PD-related phenotypes exhibited by *DJ-1ß* mutants, such as reduced locomotor activity as well as elevated ROS and protein carbonylation levels (Casani et al., 2013; Lavara-Culebras et al., 2010; Lavara-Culebras & Paricio, 2007). Our results showed that PD model flies treated with CDN1163 during development and 5 days after eclosion displayed a significant improvement of their locomotor activity when compared to individuals treated with vehicle (DMSO) (Fig. 4A). A significant reduction in protein carbonylation levels was also observed in *DJ-1ß* mutant flies treated with CDN1163 (Fig. 4B). However, no differences in H_2_O_2_ levels, a component of the total ROS pool (Casani et al., 2013; Sanz et al., 2017), were detected (Fig. 4C). To determine whether the suppression of those phenotypes was due to SERCA activation, we tested the activity of this protein in *DJ-1ß* mutant flies treated with CDN1163 by using the specific protocol mentioned above. Our results showed that SERCA activity was significantly increased in those flies when compared to vehicle-treated flies (Fig. 4D), thus indicating that PD-related phenotypes in *DJ-1ß* mutants could be caused in part by dysregulation of Ca^2+^ homeostasis. Although *Drosophila* is a valuable organism that allows to test the effect of compounds in living organisms, those shown to be beneficial in flies should be validated in mammalian models (Solana-Manrique et al., 2019). Thus, to determine the translatability of the results obtained in PD model flies, we performed cell viability assays in *DJ-1-*deficient cells pretreated with five concentrations of CDN1163 (in a range 1-50 μM) in OS conditions (see Materials and Methods section). Our results showed that treatment with CDN1163 exerted a protective effect against OS-induced cell death in *DJ-1*-deficient cells in a dosage-dependent manner (Fig. 5A). Subsequently, we performed the SERCA activity assay in cells pretreated with 5 μM CDN1163, the most effective concentration in suppressing OS-induced cell death, to determine if SERCA activity was increased in treated cells as shown in the *Drosophila* PD model. We found that cells pretreated with CDN1163 also exhibited a significant increase in SERCA activity compared to vehicle-treated cells (Fig. 5B). In summary, our results confirmed that CDN1163 exerts a protective role in PD models based on *DJ-1* deficiency through SERCA inactivation. These results indicate that CDN1163 could be a potential therapeutic compound for PD, and confirms SERCA as a therapeutic target for pharmacological intervention in PD patients.

**Figure 4.**
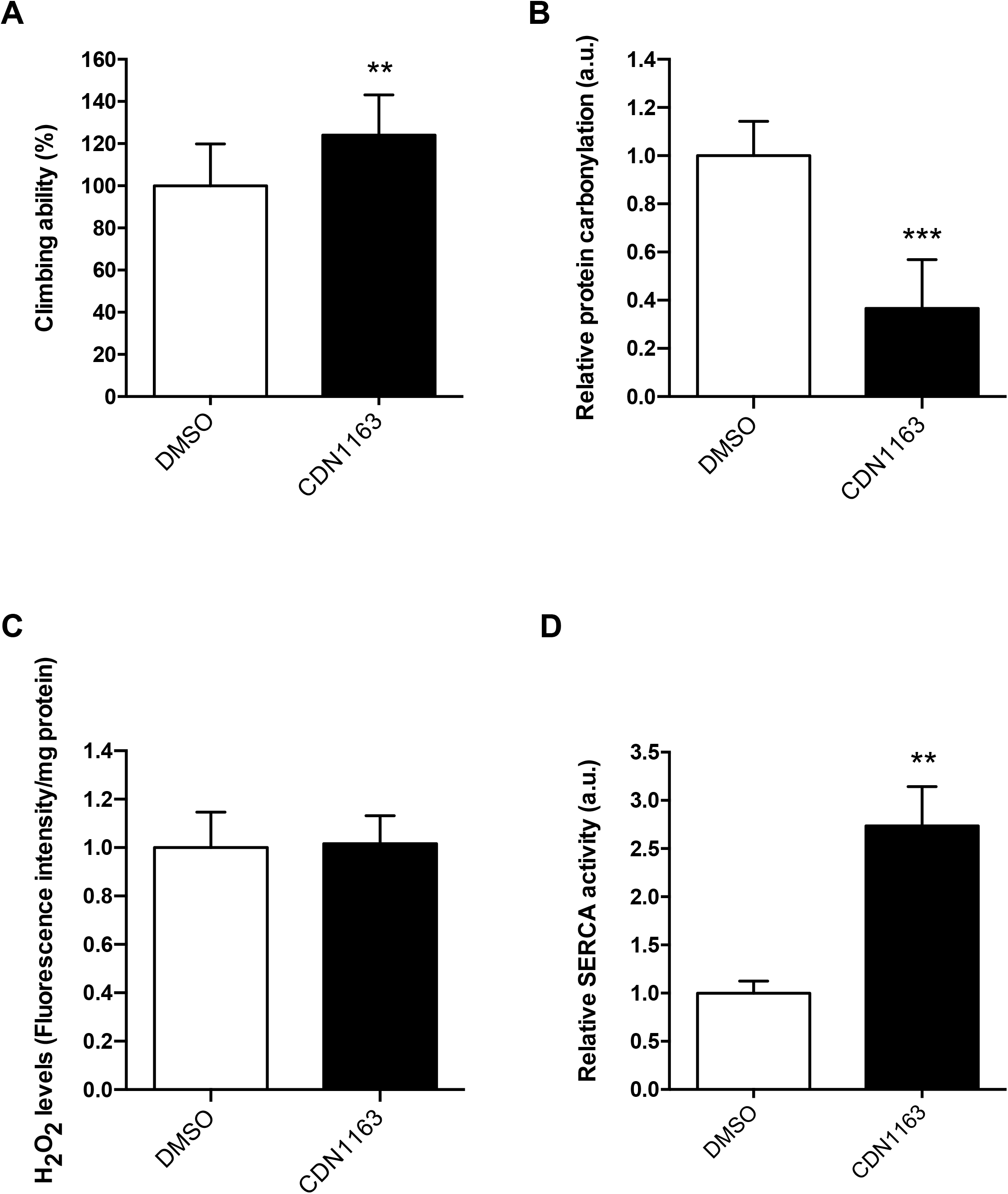
Effect of CDN1163 treatments on phenotypes exhibited by *DJ-1β* mutant flies. (A) Motor performance of *DJ-1β* mutant flies treated with 50 μM CDN1163 was analyzed using climbing assays. (B) Levels of H_2_O_2_ production in *DJ-1β* mutant flies treated with 50 μM CDN1163 were analyzed by using the *Amplex H_2_O_2_ Red Kit* (Invitrogen). Data were expressed as arbitrary units (a.u.) per mg of proteins. (C) Protein carbonylation levels in *DJ-1β* mutant flies treated with 50 μM CDN1163 were analyzed by absorbance. (D) SERCA activity was measured in 15-day-old *DJ-1β* mutant flies cultured with 50 μM CDN1163. In all cases, results were referred to data obtained in flies cultured in vehicle medium (DMSO). Error bars show s.d. from three replicates and three independent experiments (*, *P* < 0.05; **, *P* < 0.01; ***, *P* < 0.001).

**Figure 5.**
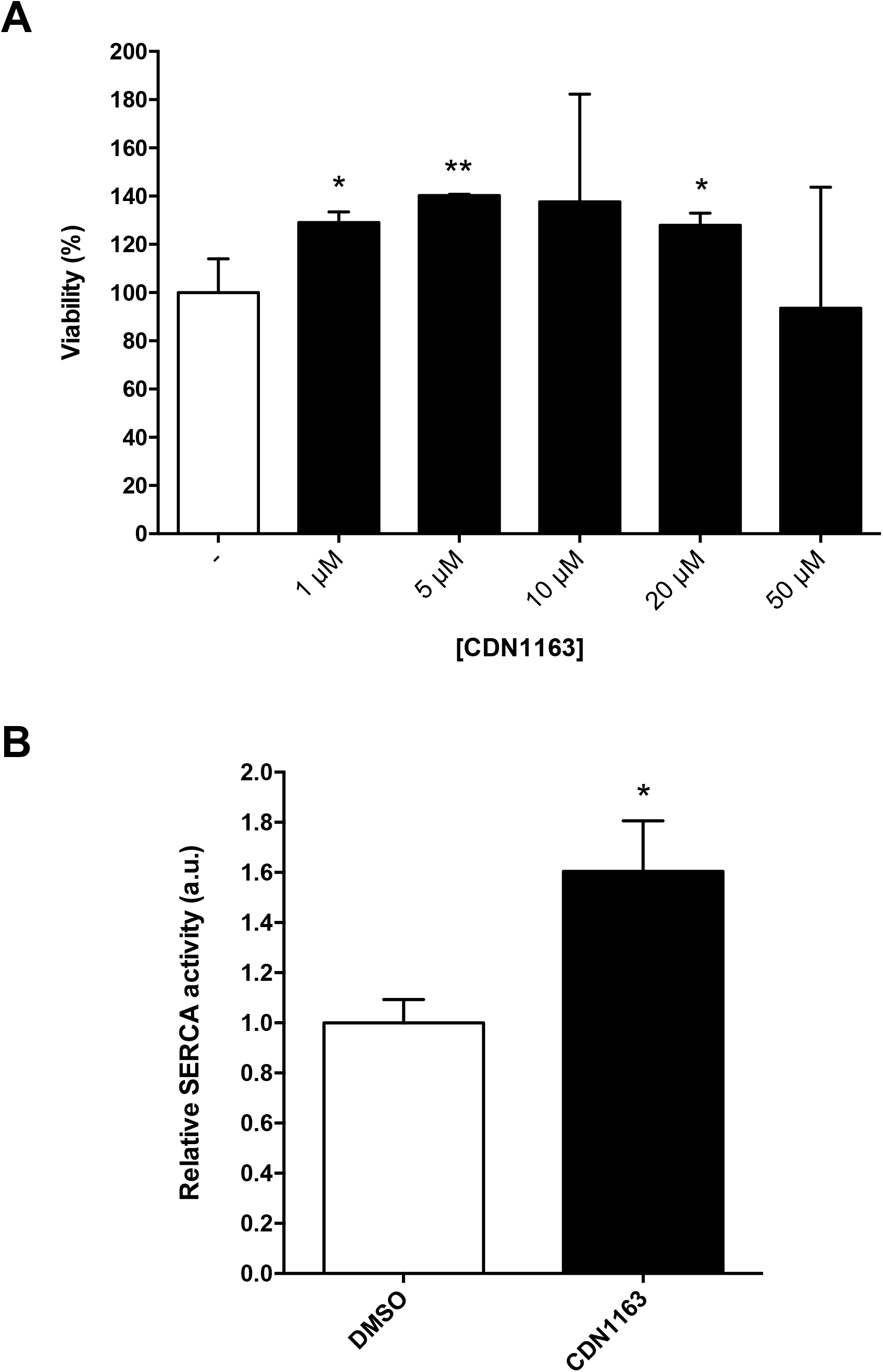
Effect of CDN1163 treatments in viability and SERCA activity in *DJ-1-*deficient cells. (A) Dosage-dependent effect of SERCA in OS-induced cell death. Viability of *DJ-1*-deficient cells was measured by MTT assays in presence of OS (induced with 100 μM H_2_O_2_). Cells were either cultured in vehicle medium (DMSO) or treated with five different concentrations of CDN1163. Results were relativized to data obtained from *DJ-1*-deficient cells cultured in vehicle (−) in presence of OS conditions. (B) SERCA activity in *DJ-1*-deficient cells treated with 5 μM CDN1163 in presence of OS conditions. Results were relativized to data from untreated *DJ-1*-deficient cells cultured in vehicle (DMSO). In all cases, error bars show s.d. from three replicates and three independent experiments (*, *P* < 0.05; **, *P* < 0.01; ***, *P* < 0.001).

## DISCUSSION

In order to obtain a better understanding of how *DJ-1* may contribute to PD pathophysiology, we carried out a redox proteomic assay in a *Drosophila* PD model based on *DJ-1β* inactivation. This study led us to identify oxidatively modified proteins in *DJ-1β* mutants whose function could be altered in such flies. We specifically focused on the SERCA calcium pump, a protein that plays a central role in Ca^2+^ homeostasis (Britzolaki et al., 2020). The SERCA protein showed increased carbonylation levels and impaired activity in *DJ-1β* mutants compared to control flies. These results were confirmed in *DJ-1*-deficient SH-SY5Y neuron-like cells. Thus, we concluded that loss of *DJ-1* function has a direct impact on the activity of this Ca^2+^ pump. In addition, SERCA activation using CDN1163 was shown to rescue PD-related phenotypes in both models. Taken together, our results demonstrated that SERCA dysfunction could be relevant for PD pathophysiology, thereby constituting a convincing therapeutic target for this disease, and that CDN1163 might play an important therapeutic role in PD.

To our knowledge, this is the first redox proteomic assay performed in a *Drosophila* PD model. OS-induced carbonyl modification of proteins has several consequences either affecting their function (activating or inhibiting them) or even promoting its elimination (Hecker & Wagner, 2018). We have recently demonstrated that the glycolytic enzymes Eno and Pfk, also identified in our redox proteomics assay (see Table 1), showed increased activity in the *Drosophila* and human cell PD models based on *DJ-1* deficiency (Solana-Manrique et al., 2020). In addition, among the proteins showing increased carbonylation levels in *DJ-1β* mutants we found actin, which is one of the most susceptible proteins to this modification under OS conditions (Castro et al., 2013). In the present study, we have focused on the SERCA protein, a Ca^2+^ pump located in the ER and ubiquitously expressed in eukaryotic cells (Primeau et al., 2018). SERCA pumps are a group of proteins that regulate cytoplasmic Ca^2+^ levels by facilitating the transport of Ca^2+^ into the ER, the most important Ca^2+^-storage organelle (Britzolaki et al., 2020; Lam & Galione, 2013; Rahate et al., 2020). We demonstrated that SERCA activity is impaired in *DJ-1*-deficient flies and human cells probably due to its carbonyl modification. This modification could be the consequence of increased ROS production in absence of *DJ-1* function, as shown in *DJ-1β* mutant flies (Casani et al., 2013; Lavara-Culebras et al., 2010). However, previous studies have demonstrated that SERCA function can be also inhibited through carbonylation by methylglyoxal (Žižková et al., 2018), a by-product of glycolysis shown to accumulate when the glycolytic pathway is overactivated (Allaman et al., 2015). Considering that glycolysis is enhanced in PD models based on *DJ-1* deficiency, in order to counteract mitochondrial dysfunction and energetic deficiency (Requejo-Aguilar et al., 2015; Solana-Manrique et al., 2020), high levels of SERCA carbonylation in the models could be also due to methylglyoxal accumulation. In any case, SERCA dysfunction may lead to an increase in cytoplasmic Ca^2+^ levels thus promoting ER stress-induced apoptosis (Lin et al., 2008). Consistently, increased [Ca^2+^] cytoplasmic levels were detected in skeletal muscle cells of *DJ-1* null mice (Shtifman et al., 2011). It has been shown that Ca^2+^ also plays an important role in coordinating organelle networks, such as mitochondria associated membranes (MAM), a major point of Ca^2+^ entrance from the ER to the mitochondria (Chernorudskiy & Zito, 2017; Erpapazoglou et al., 2017; Zaichick et al., 2017). This entrance is mediated by a protein complex composed of VDAC, IP_3_R and Grp75 (Erpapazoglou et al., 2017). Moreover, it has been demonstrated that DJ-1 directly interacts with that protein complex, probably leading to its assembly (Basso et al., 2020; Liu et al., 2019). Besides, *DJ-1* deficiency was shown to produce a decrease in MAMs number (Basso et al., 2020; Liu et al., 2019). Interestingly, we found that VDAC carbonylation levels were also increased in *DJ-1β* mutant flies in the redox proteomics assay (see Table 1). Therefore, loss of *DJ-1* function might lead to an impairment of Ca^2+^ influx from the ER to the mitochondria through MAMs either by interfering with the formation of the protein complex or by leading oxidative modification of VDAC. However, further studies will be required to confirm these hypotheses. Taken together, our results suggest that high OS levels may directly lead to alteration in Ca^2+^ homeostasis in PD models based on *DJ-1* deficiency, probably as a consequence of oxidative modification of proteins essential for this process.

Considering the impact of impaired Ca^2+^ homeostasis in PD, compounds addressed to reduce cytoplasmic Ca^2+^ have been suggested to be potential candidate therapeutics (Zaichick et al., 2017). In this study, we have tested the efficacy of CDN1163, a specific SERCA activator, in suppressing phenotypes in PD models based on *DJ-1* deficiency. Our work demonstrates that CDN116 has beneficial effects in both *Drosophila* and human cell familial PD models, thus supporting a previous work in which CDN1163 improved motor defects in 6-OHDA-lesioned rats, an idiopathic PD model (Dahl, 2017). Defects in SERCA activity are not only found in PD. AD, diabetes, muscular dystrophies or aging are pathologies in which SERCA activity is also decreased. Since these diseases are also characterized by elevated ROS levels, it is very likely that OS-induced carbonyl modification might be responsible of SERCA dysfunction (Qaisar et al., 2019; Rahate et al., 2020). Indeed, pharmacological restoration of SERCA with CDN1163 prevents OS-mediated muscle impairment in Sod1^-^ mice (Qaisar et al., 2019). In this context, the consequences of SERCA activation have been also evaluated in several models of these diseases. For instance, CDN1163 lowered fasting blood glucose and enhanced glucose tolerance in a mouse model of type 2 diabetes (Kang et al., 2016). Moreover, this compound also increased viability of rat neuronal cells and improved memory and motor performance in a mouse AD model (Krajnak & Dahl, 2018). Besides, BGP-15, a pharmacological inducer of Hsp72, increased SERCA activity under conditions of cellular stress and was found to be effective in Duchenne muscular dystrophy (Gehrig et al., 2012). Finally, overexpression of SERCA2b in the liver of obese mice was able to reduce ER stress, one of PD-related phenotypes (Park et al., 2010). Taken together, our results and those obtained in previous studies support the hypothesis that targeting Ca^2+^ dyshomeostasis, and more specifically activating SERCA activity, is a powerful therapeutic approach for PD as well as for other neurodegenerative diseases characterized by increased OS.

In summary, we demonstrated that SERCA activity is reduced in *DJ-1* mutant flies and human cells as a consequence of OS-induced carbonylation and, accordingly, treatments with the specific SERCA activator CDN1163 might have beneficial effects in PD. Hence, this supports the idea of using drugs addressed to restore Ca^2+^ homeostasis as pharmacological therapies for PD. It should be mentioned that an over-oxidized and inactive form of the DJ-1 protein has been found in brains of idiopathic PD patients (Repici & Giorgini, 2019). Therefore, it would be interesting to determine whether SERCA activity is not only altered by oxidative modification in PD models based on *DJ-1* deficiency but also in other models of familial and idiopathic PD.

## Abbreviations

AD: Alzheimer’s disease
ER: endoplasmic reticulum
MAM: mitochondrial-associated
ER: membranes
MS: mass spectrometry
OS: oxidative stress
PD: Parkinson’s disease
RyR: Ryanodine Receptors
ROS: reactive oxygen species
SERCA: sarco/endoplasmic reticulum Ca^2+^-ATPase

## ACKNOWLEDGMENTS

We are grateful to Dr. Jongkyeong Chung, the Bloomington *Drosophila* Stock Center and the Vienna *Drosophila* Research Center for providing fly stocks. The CaF2-5D2 monoclonal antibody developed by Johns Hopkins School of Medicine was obtained from the Developmental Studies Hybridoma Bank, created by the NICHD of the NIH and maintained at The University of Iowa, Department of Biology, Iowa City, IA 52242.

## SUPPLEMENTARY FIGURE

**Figure S1.**
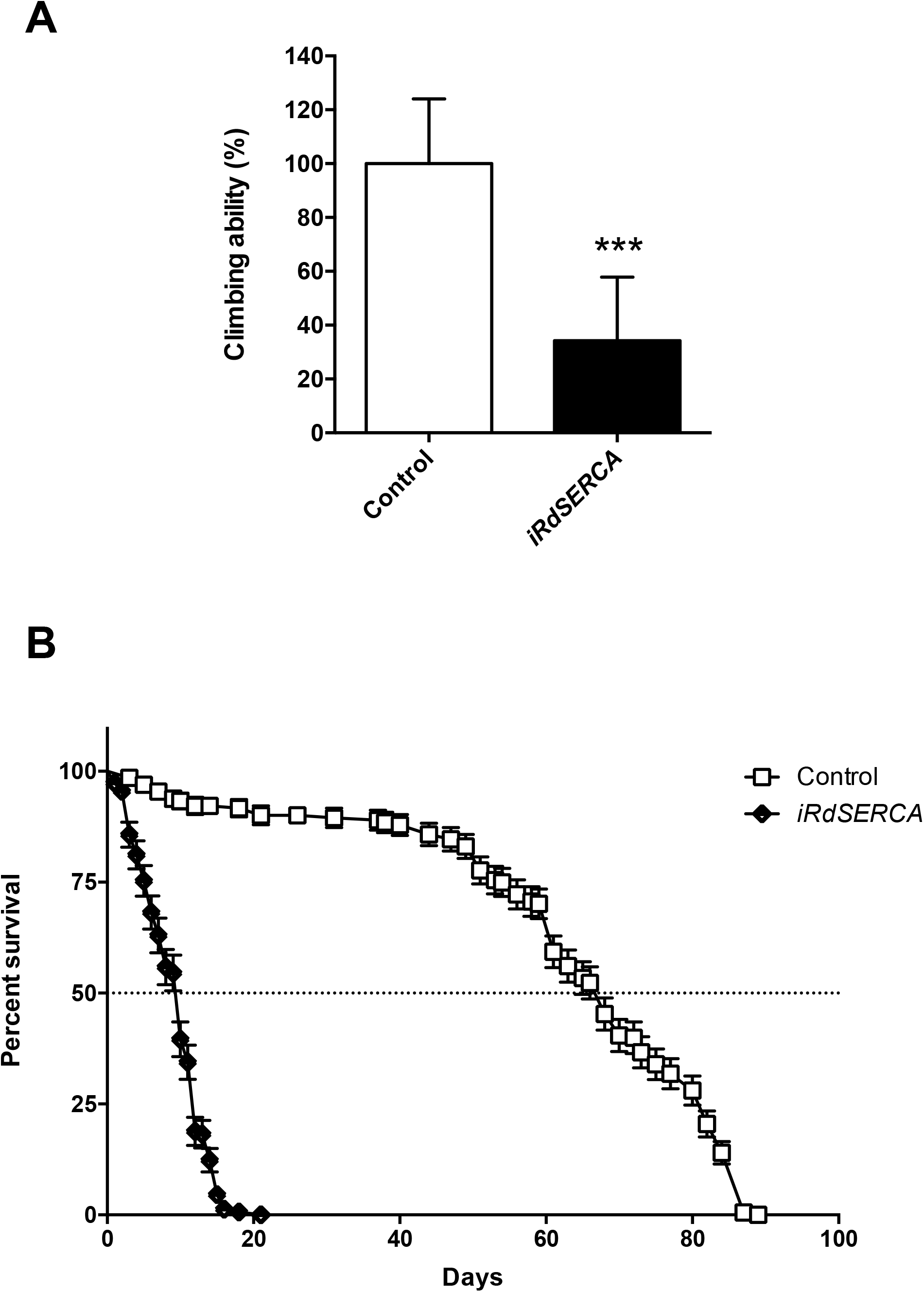
Analysis of locomotor performance and lifespan in *iRdSERCA* mutant flies. (A) Motor performance of *iRdSERCA* mutant flies was analyzed using climbing assays. Error bars show s.d. from three independent experiments (***, *P* < 0.001). (B) Comparison of lifespan curves of *iRdSERCA* mutant (white squares) and *arm*-GAL4/+ control flies (black diamond). The median and maximum lifespan values are as follows: 68 and 89 days for *arm-GAL4/+* controls; 10 and 21 days for *iRdSERCA* mutant flies. The significance of the difference between survival curves was analyzed using Kaplan-Meier log-rank statistical test (****, *P* < 0.0001).

**Table S1.** Functional role of the highly carbonylated proteins identified in the redox proteomics assay.

| Spot number | Protein | Functional role |
| --- | --- | --- |
| 175 | V-type proton ATPase catalytic subunit A isoform 2 | Catalytic subunit of the peripheral VI complex of vacuolar ATPase (V-ATPase). V-ATPase is responsible of acidifying a variety of intracellllular compartemts in eukaryotic cells |
| 336 | Tyrosyl-tRNA synthethase | Exhibits tyrosine-tRNA ligase activity, catalyzes the ligation of tyrosine to its cognate tRNAs in the cytoplasm |
| 384 | Myosin heavy chain, muscle | Muscle contraction |
| 496 | Enolase | Exhibits phosphopyruvate hydratase activity. Enolase is involved in glucose homeostasis |
| 555 | Probable cytrate synthase, mitochondrial | Plays a role in controlling neuronal activity and seizure susceptibility |
|  | ATP-dependent 6-phosphofructokinase | Catalyzes the phosphorylation of D-fructose 6-phosphate to fructose 1,6-bisphosphate by ATP, the first committing step of glycolysis |
| 625 | Vitellogenin-3 | Vitellogenin is the major yolk protein of eggs where it is used as a food source during embryogenesis |
| 716 | Actin | Actin are highly conserved proteins that are involved in various types of cells motility and are ubiquitously expressed in all eukaryotic cells. Multiple isoforms are involved in various cellular functions such as cytoeskeleton structure, cell motility, chromosome movement and muscle contraction |
| 731 | ATP-dependent 6-phosphofructokinase | Catalyzes the phosphorylation of D-fructose 6-phosphate to fructose 1,6-bisphosphate by ATP, the first committing step of glycolysis |
| 795 | Vitellogenin-2 | Vitellogenin is the major yolk protein of eggs where it is used as a food source during embryogenesis |
| 971 | Voltage-dependent anion-selective cannell | Forms a channel through the cell membrane that allows diffusion of small hydrophilic molecules |
| 1061 | Arginine kinase | Exhibits creatine kinase activity and transfers a phosphate group, usually from ATP, to creatine substrate |
| 1310 | Calcium-transporting ATPase sarcoplasmic | This magnesium-dependent enzyme catalyzes the hydrolysis of ATP coupled with the transport of calcium to the endoplasmic reticulum |

